# Draft Genome Sequence of *Pedococcus* sp. Strain 5OH_020 Isolated from California Grassland Soil

**DOI:** 10.1101/2023.01.13.523987

**Authors:** Myka A. Green, Zoila I. Alvarez-Aponte, Valentine V. Trotter, Michiko E. Taga

## Abstract

The draft genome sequence of the soil bacterium *Pedococcus* sp. 5OH_020, isolated on a natural vitamin B_12_ analog, contains 4.4 Mbp with 4,108 protein-coding genes. Its genome encodes B_12_-dependent enzymes including methionine synthase and class II ribonucleotide reductase. Taxonomic analysis suggests it is a novel species within the genus *Pedococcus*.

## Announcement

*Pedococcus* sp. strain 5OH_020 was isolated from ungrazed grassland topsoil at the Hopland Research and Extension Center (39.00 N 123.08 W) by limiting dilution in methionine-dropout VL60 medium containing 0.1 g/L each of xylan, glucose, xylose, and N-acetyl glucosamine (1), amended with 10 nM 5-hydroxybenzimidazolyl cobamide ([5-OHBza]Cba) that we synthesized according to (2). [5-OHBza]Cba is a commercially-unavailable cobamide cofactor typically biosynthesized by methanogens and found in diverse environments such as animal gastrointestinal tracts and contaminated groundwater (3, 4, 5). To our knowledge, this is the first report of a bacterium isolated on a cobamide other than vitamin B_12_ (cobalamin).

Strain 5OH_020 was cultured at 28 ºC on 2X VL60 agar, or liquid VL60 medium, amended with 10 nM [5-OHBza]Cba. The isolate is aerobic, Gram-positive, and cocci-shaped (Figure 1 inset), forming circular, cream-colored colonies after approximately 12 days and requires [5-OHBza]Cba or B_12_ for growth. We performed whole genome sequencing after noting low similarity to its closest 16S rRNA pairwise matches (<98.65%) (6).

**Figure 1.**
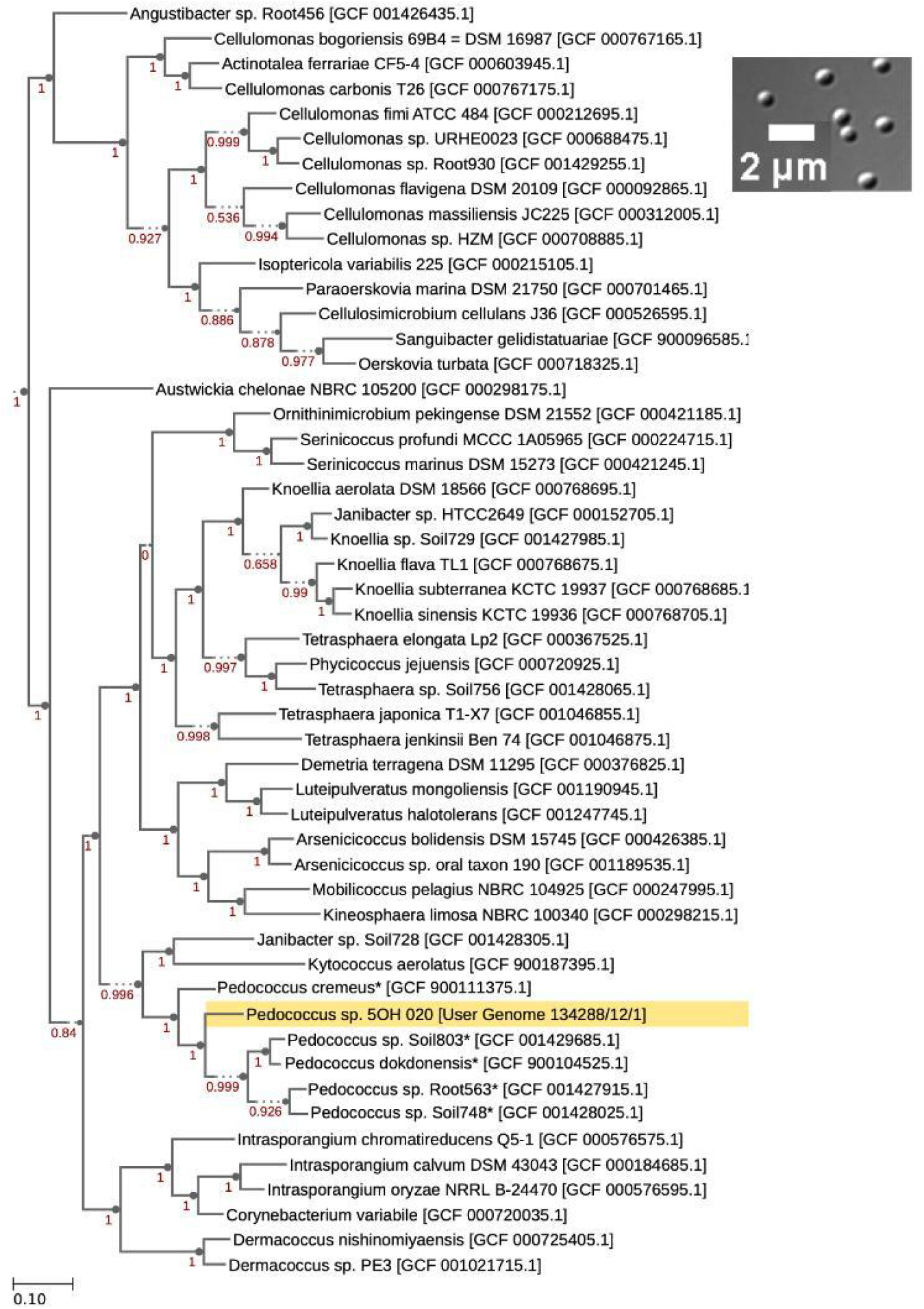
Morphology and phylogeny of *Pedococcus* sp. strain 5OH_20. Insert Genome into Species Tree v2.2.0 (16) application with parameters set to 50 neighboring species was used to generate a phylogenetic tree of representative members of class Actinomycetia. Strain 5OH_020 is highlighted in yellow. The species marked with * were corrected from the genus *Phycicoccus* to *Pedococcus* to match current taxonomic classifications according to the NCBI database. Numbers in red represent bootstrap values. (Inset) Differential interference contrast (DIC) microscopy at 100x magnification with a Zeiss AxioImager DIF M1 microscope. Image captured by Hamamatsu Orca 03 camera.

DNA was extracted using the DNeasy Blood and Tissue Kit (Qiagen) protocol for Gram-positive bacteria. Subsequent library preparation and sequencing steps were carried out by Novogene Bioinformatics Technology Co., Ltd (Beijing, China). Library construction was performed using NEBNextⓇ DNA Library Prep Kit following the manufacturer’s instructions. After end repairing, dA-tailing, and further ligation with NEBNext adapter, DNA was sequenced using the Illumina Novaseq 6000 platform (Illumina Inc., San Diego, CA, USA) to generate 150 bp reads, yielding 9,600,674 total reads following a quality control step.

Data processing and analysis were performed with the Department of Energy Systems Biology Knowledgebase (KBase) (7) using default parameters unless otherwise noted. Raw reads were trimmed and further quality controlled using Trimmomatic v0.36 (8). The genome was assembled *de novo* via SPAdes v3.15.3 (9) and quality checked with QUAST v4.4 (10), generating the assembly statistics in Table 1. The genome was annotated and re-annotated with Prokka v.1.14.5 (11) and distilled with DRAM v0.1.2 (12). The 4.4 Mbp genome was quality checked using CheckM v1.0.18 (13).

**Table 1.** Assembly statistics generated by QUAST v4.4.

| Assembly Statistic | Result |
| --- | --- |
| Number of contigs | 33 |
| Largest contig (bp) | 783 609 |
| Total length (bp) | 4 427 096 |
| Coverage | 325x |
| N50 (bp) | 428 675 |
| N75 (bp) | 278 038 |
| L50 | 5 |
| L75 | 8 |
| G+C content (%) | 69.75 |

The genome was classified as genus *Pedococcus* in the Actinobacteria phylum, with no further classification at the species level by GTDB-Tk v1.7.0 (14). The amino acid identity with *Pedococcus* sp. soil748 was 95.61%. FastANI v1.33 indicates 81.17% average nucleotide identity with the genus type strain *Pedococcus dokdonensis* (15). *Pedococcus* sp. strain 5OH_020 forms an independent branch within the *Pedococcus* clade (Figure 1) (16).

*Pedococcus* sp. 5OH_020 belongs to class Actinomycetia, which is renowned for its production of clinically useful natural products (17). Among the 4,108 annotated protein-coding genes, two natural product biosynthetic clusters were detected by antiSMASH v6.0 (18). The genome also encodes enzymes predicted to degrade arabinan, xyloglucan, and amorphous cellulose, suggesting 5OH_020 may be capable of degrading plant matter present in soil.

Several cobalamin-dependent enzymes including methionine synthase and class II ribonucleotide reductase are encoded in the genome. No cobamide biosynthesis genes were present, consistent with the cobamide dependence observed in culture. Thus, it likely relies on other microbes in the soil environment to provide cobamides.

## Data Availability

The draft genome sequence of *Pedococcus* sp. strain 5OH_020 was deposited in DDBJ/ENA/GenBank under the accession <u>JAPYZ000000000</u>. The version described in this paper is version JAPYZV010000000. The NCBI SRA accession for the raw reads is <u>SRR22578002</u>. The BioProject and BioSample accessions for this project are <u>PRJNA891672</u> and <u>SAMN31357841</u> respectively.

## Acknowledgements

We thank Dr. Denise Schichnes and the CNR Biological Imaging Facility at the University of California, Berkeley, for microscopy guidance, and Dr. Hans Carlson and Dr. Adam Deutschbauer for their advice on whole genome sequencing. This research was supported by the Sponsored Projects for Undergraduate Research (SPUR) program at UC Berkeley (M.A.G.) and the US Department of Energy grant DE-SC0020155 to M.E.T. Z.I.A.A. was supported, in part, by a Grace Kase Fellowship from the Kase/Tsujimoto Foundation and a GEM Fellowship.

